# Overlapping coactivator function is required for transcriptional activation by the *Candida glabrata* Pdr1 transcription factor

**DOI:** 10.1101/2024.05.24.595833

**Authors:** Thomas P. Conway, Lucia Simonicova, W. Scott Moye-Rowley

## Abstract

Azole resistance in the pathogenic yeast *Candida glabrata* is a serious clinical complication and increasing in frequency. The majority of resistant organisms have been found to contain a substitution mutation in the Zn2Cys6 zinc cluster-containing transcription factor Pdr1. These mutations typically lead to this factor driving high, constitutive expression of target genes like the ATP-binding cassette transporter-encoding gene *CDR1*. Overexpression of Cdr1 is required for the observed elevated fluconazole resistance exhibited by strains containing one of these hyperactive *PDR1* alleles. While the identity of hyperactive *PDR1* alleles has been extensively documented, the mechanisms underlying how these gain-of-function (GOF) forms of Pdr1 lead to elevated target gene transcription are not well understood. We have used a tandem affinity purification (TAP)-tagged form of Pdr1 to identify coactivator proteins that biochemically purify with the wild-type and two different GOF forms of Pdr1. Three coactivator proteins were found to associate with Pdr1: the SWI/SNF complex Snf2 chromatin remodeling protein and two different components of the SAGA complex, Spt7 and Ngg1. We found that deletion mutants lacking either *SNF2* or *SPT7* exhibited growth defects, even in the absence of fluconazole challenge. To overcome these issues, we employed a conditional degradation system to acutely deplete these coactivators and determined that loss of either coactivator complex, SWI/SNF or SAGA, caused defects in Pdr1-dependent transcription. A double degron strain that could be depleted for both SWI/SNF and SAGA exhibited a profound defect in *PDR1* autoregulation, revealing that these complexes work together to ensure high level Pdr1-dependent gene transcription.

## Introduction

Treatment options for fungal infections are primarily limited to three classes of antifungal drugs, azoles, echinocandins, and polyenes, with the azole drug, fluconazole, being the most commonly prescribed therapeutic agent for fungal infections worldwide (Nett and Andes 2016; Krysan 2017). Azole drugs, like fluconazole, inhibit growth of fungal pathogens by inhibiting a key enzyme in the ergosterol biosynthesis pathway, lanosterol α-14 demethylase (Sagatova *et al*. 2015). The incidence of azole resistance among clinical *C*. *glabrata* isolates is considered so high that drug resistant *Candida* species have been designated as a critical threat by the World Health Organization (Fisher and Denning 2023). Azole resistance in *C. glabrata* has been linked primarily to nonsynonymous point mutations in the open reading frame of the transcription factor-encoding gene, *PDR1*, which result in hyperactive, gain-of-function (GOF) Pdr1 variants (Ferrari *et al*. 2009). Pdr1 activates its own expression via positive transcriptional autoregulation, with this induced expression of Pdr1 being essential for the subsequent activation of a battery of genes required for azole resistance, including the ATP-binding cassette (ABC) multi-drug transporter, *CDR1* (Sanglard *et al*. 2001; Healey and Perlin 2018). Multiple groups have shown that the overexpression of Cdr1 is a direct consequence of constitutively active Pdr1 forms (Vermitsky *et al*. 2006; Ferrari *et al*. 2009; Tsai *et al*. 2010). Determining how wild-type and GOF variants of Pdr1 induce and maintain gene activation under azole stress is critical to the prospect of enhancing therapeutic options as the incidence of azole resistant isolates increases.

Pdr1 is a Zn_2_Cys_6_ transcription factor (Vermitsky and Edlind 2004) (Tsai *et al*. 2006; Paul *et al*. 2011; Khakhina *et al*. 2018), with its DNA-binding domain (DBD) contained within the first 300 amino acids at the N-terminus of the protein. The region containing the DBD precedes a central regulatory domain (CRD) comprised of approximately 700 amino acids, which is in turn followed by a C-terminal transactivation domain (TAD). It has been demonstrated that the TAD of Pdr1 interacts directly with Med15A, a subunit of the Mediator transcriptional complex (reviewed in (Kornberg 2005; Casamassimi and Napoli 2007)), and that this interaction is required for Pdr1 to activate gene expression (Thakur *et al*. 2008). The Mediator complex serves as a link between sequence-specific binding of transcription factors, like Pdr1, and RNA polymerase II (recently reviewed in (Malik and Roeder 2023)). Furthermore, past work has shown that GOF mutations can occur in either the CRD or TAD of the Pdr1 protein, both resulting in constitutively active Pdr1 forms (Ferrari *et al*. 2009). We previously demonstrated that two distinct Pdr1 GOF mutant forms, R376W and D1082G, have a differential dependence on Med15A for their hyper-resistant phenotypes (Simonicova and Moye-Rowley 2020). Upon removal of Med15A, a greater increase in fluconazole susceptibility is seen for cells expressing R376W Pdr1 than for cells expressing D1082G Pdr1 (Simonicova and Moye-Rowley 2020). While a large number of GOF variants of Pdr1 have been identified (Ferrari *et al*. 2009), little is known about how these mutations alter the mechanism of Pdr1-driven gene expression.

To determine the proteins that directly interact with both wild-type and two different GOF forms of Pdr1, we utilized a tandem affinity purification (TAP)-tagged fusion approach (Puig *et al*. 2001). Wild-type, R376W, and D1082G forms of Pdr1 were amino-terminally TAP-tagged and purified biochemically. Mass spectrometry was used to identify accessory proteins associated with each form of Pdr1. We found that the SWI/SNF subunit, Snf2, and the SAGA subunits, Spt7 and Ngg1/Ada3, were co-purified with all forms of Pdr1. We provide evidence for the Pdr1-dependent localization and azole-induced recruitment of SWI/SNF and SAGA subunits to the promoters of azole-responsive genes, and we show that the level of promoter recruitment of the coactivator complexes varies depending on the Pdr1 variant expressed. Finally, we demonstrate that simultaneous depletion of both Spt7 and Snf2 blocked fluconazole activation of the *PDR1* gene but not *CDR1*, demonstrating the unique coactivator requirements of these two Pdr1-controlled target genes.

## Materials and Methods

### Strains and Growth Conditions

All strains used in this study are listed in Table 1. For non-selective growth and drug treatment, *C. glabrata* strains were grown at 30°C in YPD medium (1% yeast extract, 2% peptone, 2% glucose). Selection for transformants occurred on either YPD media supplemented with 50 µg/ml nourseothricin (Jena Bioscience, Jena, Germany) and 2 mM methionine or minimal SD media (yeast nitrogen base with ammonium sulfate) supplemented with appropriate amino acids. When applicable, transformation cassettes were recycled via growth in synthetic complete media (2% glucose, 1mM estradiol) without methionine.

**Table 1.** Strain List, List of strains used in this study.

| Strain name | Background | Genotype |
| --- | --- | --- |
| CBS138 | CBS138 | Wild-type |
| TCCG200 | CBS138 | *loxP::TAP-PDR1* |
| TCCG201 | CBS138 | *loxP::TAP-PDR1-R376W* |
| TCCG202 | CBS138 | *loxP::TAP-PDR1-D1082G* |
| TCCG3 | CBS138 | *pdr1Δ::loxP* |
| TCCG122 | CBS138 | *cdr1Δ::loxP* |
| TCCG20 | CBS138 | *ngg1Δ::natMX6* |
| TCCG12 | CBS138 | *snf2Δ::natMX6* |
| TCCG61 | CBS138 | *spt7Δ::natMX6* |
| TCCG67 | CBS138 | *PDR1::loxP NGG1-3XHA::natMX6* |
| TCCG73 | CBS138 | *pdr1Δ::loxP NGG1-3XHA::natMX6* |
| TCCG85 | CBS138 | *PDR1-R376W::loxP NGG1-3XHA::natMX6* |
| TCCG79 | CBS138 | *PDR1-D1082G::loxP NGG1-3XHA::natMX6* |
| TCCG111 | CBS138 | *PDR1::loxP SNF2-3XHA::natMX6* |
| TCCG142 | CBS138 | *pdr1Δ::loxP SNF2-3XHA::natMX6* |
| TCCG116 | CBS138 | *PDR1-R376W::loxP SNF2-3XHA::natMX6* |
| TCCG114 | CBS138 | *PDR1-D1082G::loxP SNF2-3XHA::natMX6* |
| TCCG65 | CBS138 | *PDR1::loxP SPT7-3XHA::natMX6* |
| TCCG71 | CBS138 | *pdr1Δ::loxP SPT7-3XHA::natMX6* |
| TCCG83 | CBS138 | *PDR1-R376W::loxP SPT7-3XHA::natMX6* |
| TCCG77 | CBS138 | *PDR1-D1082G::loxP SPT7-3XHA::natMX6* |
| KKY2001 | CBS138 | *his3Δ::FRT leu2Δ::FRT trp1Δ::FRT* |
| TCCG32 | KKY2001 | *trp1Δ::ScLEU2-OsTIR1-9XMyC* |
| TCCG63 | KKY2001 | *NGG1-3XminiAID-5XFLAG::his3MX6*<br>*trp1Δ::ScLEU2-OsTIR1-9XMyC* |
| TCCG163 | KKY2001 | *PDR1-R376W::loxP NGG1-3XminiAID-*<br>*5XFLAG::his3MX6 trp1Δ::ScLEU2-OsTIR1-*<br>*9XMyC* |
| TCCG161 | KKY2001 | *PDR1-D1082G::loxP NGG1-3XminiAID-*<br>*5XFLAG::his3MX6 trp1Δ::ScLEU2-OsTIR1-*<br>*9XMyC* |
| TCCG53 | KKY2001 | *SNF2-3XminiAID-5XFLAG::his3MX6*<br>*trp1Δ::ScLEU2-OsTIR1-9XMyC* |
| TCCG150 | KKY2001 | *PDR1-R376W::loxP SNF2-3XminiAID-*<br>*5XFLAG::his3MX6 trp1Δ::ScLEU2-OsTIR1-*<br>*9XMyC* |
| TCCG148 | KKY2001 | *PDR1-D1082G::loxP SNF2-3XminiAID-*<br>*5XFLAG::his3MX6 trp1Δ::ScLEU2-OsTIR1-*<br>*9XMyC* |
| TCCG55 | KKY2001 | *SPT7-3XminiAID-5XFLAG::his3MX6*<br>*trp1Δ::ScLEU2-OsTIR1-9XMyC* |
| TCCG156 | KKY2001 | *PDR1-R376W::loxP SPT7-3XminiAID-*<br>*5XFLAG::his3MX6 trp1Δ::ScLEU2-OsTIR1-*<br>*9XMyC* |
| TCCG154 | KKY2001 | *PDR1-D1082G::loxP SPT7-3XminiAID-*<br>*5XFLAG::his3MX6 trp1Δ::ScLEU2-OsTIR1-*<br>*9XMyC* |
| TCCG180 | KKY2001 | *SNF2-3XminiAID-5XFLAG::his3MX6 SPT7-3XminiAID-5XFLAG::bleMX6 trp1Δ::ScLEU2-OsTIR1-9XMyC* |

Isogenic strains carrying the wild type or gain-of-function mutations in *PDR1* allele (referred to as R376W or D1082G) were generated by integrating the respective *PDR1* allele in a markerless manner into the genome of CBS138 *pdr1*11 strain (Simonicova and Moye-Rowley 2020). The isogenic strains carrying wild type, R376W, or D1082G alleles that were used for the purification of various Pdr1 forms for the mass spectrometric analyses were constructed by the same approach as mentioned above, with the exception that the alleles were first cloned in amino-terminal fusion with tandem affinity purification (TAP)-tag. Both, the untagged and TAP-tagged sets of isogenic *PDR1* strains generated in this manner carry no other alterations to their genome.

### TAP-tag purification and Multidimensional Protein Identification Technology (MudPIT)

The protocol was modified from (Rigaut *et al*. 1999). Briefly, 500 OD_600_ units of mid-log cells grown in YPD were harvested, washed in water and resuspended in 500 μl of NP-40 lysis buffer (15 mM Na_2_HPO_4_, 10 mM NaH_2_PO4, 1% Nonidet, 150 mM NaCl, 2 mM EDTA) complemented by 1 mM PMSF, 1 mM DTT, 1X Complete protease inhibitor, 50 mM sodium fluoride and 0.1 mM sodium vanadate. Cells were lysed in the presence of glass beads in four cycles of vortexing for 5 minutes at maximum speed followed by cooling on ice for one minute. Lysates were cleared by centrifugation and the supernatant was loaded onto an IgG-sepharose column, followed by incubation for 4 hours at 4°C. Beads were washed (25 mM Tris-HCl (pH 8.0), 0.1% Nonidet, 300 mM NaCl) followed by the second wash using the same buffer containing 150 mM NaCl.

TAP-Pdr1 proteins were eluted by cleavage with TEV protease (100 U) overnight at 4°C Eluates were then applied onto the calmodulin-sepharose column in the presence of 2 mM calcium chloride, incubated for 2 hours at 4°C followed by washing in the same column buffer. Bound Pdr1 proteins were eluted from the column by addition of 20 mM EGTA. The eluates were TCA-precipitated, washed with acetone and dried before the mass-spectrometric analysis. The MudPIT analysis was performed as described in (Paul *et al*. 2018) and performed by Dr. Hayes Macdonald (Vanderbilt University Medical Center). Raw MudPIT data containing the list of all Pdr1-interacting proteins can be found in Supplemental Table 1.

### *C. glabrata* transformation and gene modification

Gene disruptions and epitope-tagged alleles were generated using a CRISPR-Cas9 method (Grahl *et al*. 2017). After electroporation, cells were plated immediately on CSM agar plates for auxotrophic selection. For nourseothricin selection, transformed cells were grown in YPD at 30°C and 200 rpm for 3 hours after electroporation then plated on YPD agar supplemented with 50 μg/ml nourseothricin. Selection plates were incubated at 30°C for 48 hours, then individual colonies were isolated and screened by PCR for targeted insertion of the transformation construct.

### Spot Test Drug Treatment

*C. glabrata* strains were grown to mid-log phase and spotted in ten-fold serial dilutions on YPD containing the indicated concentrations of fluconazole (LKT Laboratories, Ing. #F4682) and/or 1-naphthaleneaceitc acid (NAA/auxin, Sigma-Aldrich). Plates were incubated at 30°C for 24 to 48 hours prior to imaging.

### Western Blot Analysis

Proteins were extracted from 3 OD_600_ units of mid-log cells as previously described (Paul *et al*. 2011), resuspended in urea sample buffer (8 M urea, 1% 2-mercaptoethanol, 40 mM Tris-HCL pH 8.0, 5% SDS, bromophenol blue), and incubated at room temperature until all proteins had solubilized. Extract aliquots were resolved on precast ExpressPlus 4-12% gradient gel (GenScript #M41212), transferred to a nitrocellulose membrane, and blocked with 5% nonfat dry milk. For the analysis of Pdr1 and Cdr1 expression, the blocked membranes were probed with anti-Pdr1 antibody (Paul *et al*. 2014) and anti-Cdr1 antibody (Vu *et al*. 2019), respectively. An anti-FLAG antibody was used to monitor coactivator depletion where the auxin-inducible degradation strategy was employed. All membranes were also probed with 12G10 anti-alpha-tubulin mouse monoclonal antibody (Developmental Studies Hybridoma Bank, University of Iowa). The secondary Li-Cor antibodies IRD dye 680RD goat anti-rabbit (#926-68021) and IRD dye 800LT goat anti-mouse (#926-32210) were used along with a Li-Cor infrared imaging system and Image Studio Lite software (Li-Cor) for detection of blotted proteins.

### Quantification of Transcript Levels by Real Time qPCR

Total RNA was extracted from samples containing 6 OD_600_ units of mid-log phase cells using Trizol (Invitrogen #15596026) and chloroform. A RNeasy Mini Kit (Qiagen #74104) was used for RNA purification, and cDNA was generated from 0.5 micrograms of purified RNA from each sample using an iScript cDNA Synthesis Kit (Bio-Rad #1708890). qPCR was conducted using iTaq Universal SYBR Green Supermix (Bio-Rad #1725151). Target gene transcript levels were normalized to 18S rRNA transcript levels, and the 2^-ΔΔCt^ method (Livak and Schmittgen 2001) was used for calculations of fold change between samples for transcript levels of the genes of interest. All measurements represent the results of two technical replicates each from two independent biological replicates. Statistical analysis was carried out using a paired t test for within strain comparisons between different conditions and an unpaired t test was used for comparisons between strains where applicable.

### Chromatin Immunoprecipitation and qPCR

Chromatin immunoprecipitation (ChIP) was performed as previously described (Paul *et al*. 2014; Vu *et al*. 2021). Samples containing 75 OD_600_ units of mid-log phase cells were fixed and DNA-protein interactions crosslinked by treatment with 1% formaldehyde. The crosslinking reaction was subsequently inhibited by the addition of glycine and cells were washed with 1X PBS. Cell pellets were resuspended in FA-lysis buffer (50 mM HEPES-KOH, 140 mM NaCl, 1 mM EDTA, 1% Triton X-100, 0.1% sodium deoxycholate, supplemented with 1 mM PMSF, and 1X complete protease inhibitor). Approximately 1 ml of 0.5 mm glass beads were added to the cell suspension and the samples were vortexed at 4°C for a total of 10 minutes in 2 minute intervals separated by a 1 minute pause on ice. The resulting lysate was separated from the glass beads and 130 µl from each sample was added to an AFA Fiber Pre-Slit Snap-Cap microtube (Covaris). Chromatin was sheared in a E220 Focused-Ultrasonicator using the following conditions: peak incident power (W): 175, duty factor: 20%, cycles per burst: 200, treatment time (seconds): 720, temperature (°C): 7, sample volume (µl): 130, under the presence of an E220 intensifier (pn500141). Debris was removed from each sample by centrifugation and a portion of the supernatant was separated for use as an input control. Immunoprecipitation was performed using an anti-HA monoclonal antibody (2-2.214; Invitrogen) against the 3X HA-tagged coactivators. The antibody was added to the lysate (1:50) and incubated at 4°C for 2 hours. Dynabeads Protein G magnetic beads (Invitrogen) were added, and the mixture was incubated overnight at 4°C on a nutator. The remaining wash steps and reversal of the crosslink were carried out as previously described (Vu *et al*. 2021).

## Results

### Identification of Pdr1 protein partners

Positive regulation of gene expression by the transcription factor Pdr1 is critical for the response to azole stress in *C. glabrata*. Previously, we have identified and characterized several potential regulators of Pdr1 activity by expressing a tandem affinity purification (TAP)-tagged form of wild-type Pdr1 in wild-type cells of *C. glabrata* or cells that lack a functional mitochondrial genome (ρ^0^ cells) (Paul and Moye-Rowley 2014; Paul *et al*. 2018). While Pdr1 activity is increased in mitochondrial mutants, the primary mechanism of Pdr1 activation in fluconazole resistant clinical isolates is the acquisition of specific nonsynonymous point mutations in the open reading frame of *PDR1* that result in constitutively active, gain-of-function (GOF) alleles (Ferrari *et al*. 2009). Our earlier data demonstrated that different GOF Pdr1 variants showed variable dependence on coactivators required for their GOF phenotype (Simonicova and Moye-Rowley 2020). Currently, the only Pdr1 coactivator characterized in detail is the Mediator subunit Med15A (Thakur *et al*. 2008; Paul *et al*. 2011).

To identify Pdr1 protein partners, we analyzed three isogenic strains that varied only in the *PDR1* allele they expressed. The different *PDR1* alleles corresponded to wild-type Pdr1 or one of two GOF Pdr1 forms, R376W Pdr1 and D1082G Pdr1, which are of clinical relevance and have been demonstrated to have different requirements for maintaining their constitutively active state (Ferrari *et al*. 2009; Simonicova and Moye-Rowley 2020), Figure 1A). We TAP-tagged the amino-terminus of each *PDR1* allele and integrated them into the genome of the wild-type *C. glabrata* strain CBS138 (Figure 1A). To validate their function, TAP-tagged Pdr1 strains were analyzed by spot test assay for susceptibility to fluconazole and compared to strains expressing the corresponding untagged Pdr1 variants. Additionally, Pdr1 and Cdr1 protein levels were assayed by western blot (Figure 1B). All TAP-tagged strains exhibited susceptibility to fluconazole comparable to the corresponding untagged strains (Figure 1B) and both TAP-tagged Pdr1-GOFs drove higher expression of Cdr1 than TAP-tagged wild-type Pdr1 (Figure 1C).

**Figure 1:**
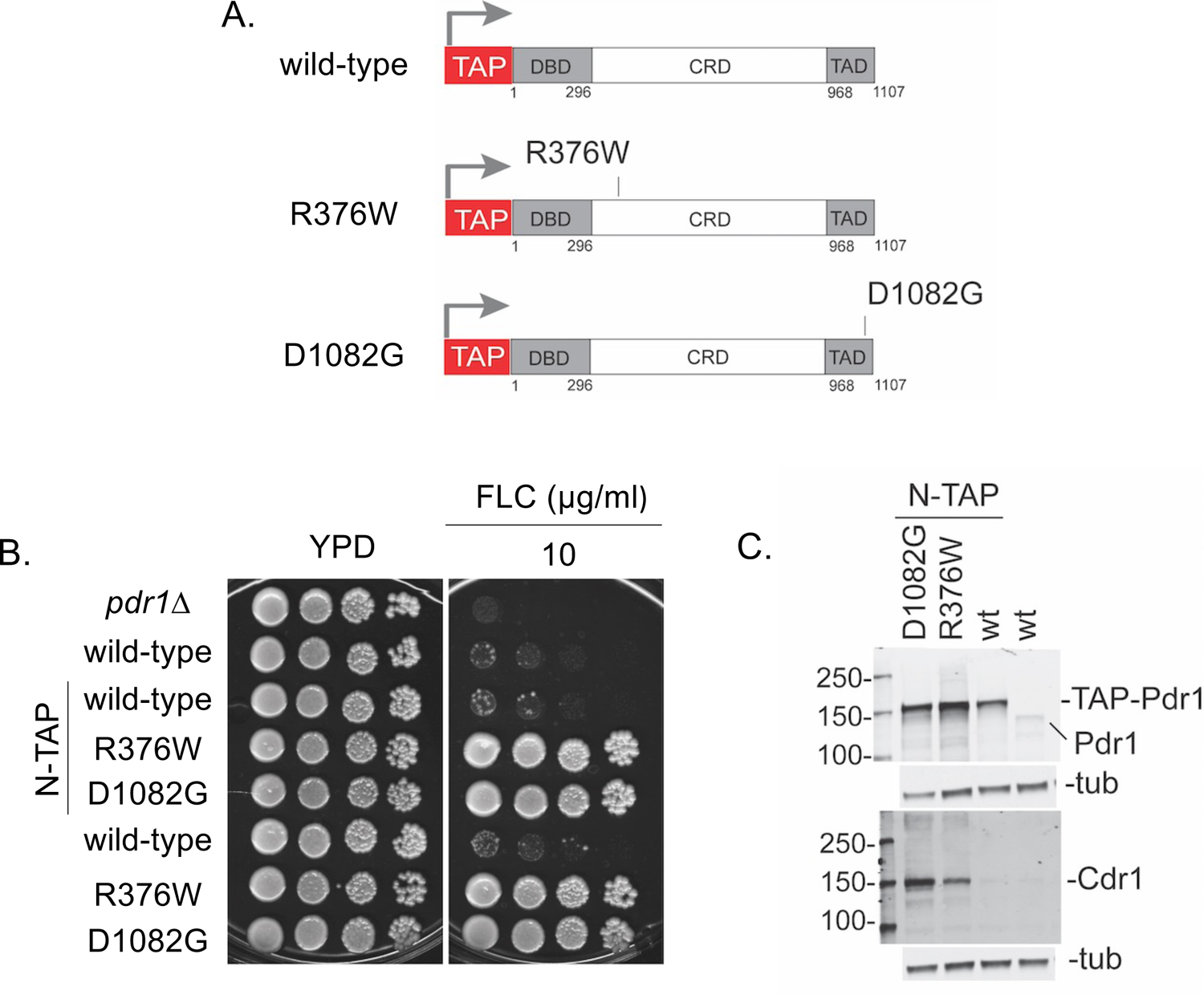
Phenotypic characterization of Pdr1 gain-of-function mutations in amino-terminal fusion with TAP-tag. A. Functional domains of Pdr1 (DBD, DNA binding domain; CRD, central regulatory domain and TAD, transactivation domain) with the location of gain-of-function mutations indicated. B. Strains expressing Pdr1 alleles (i.e. wild-type, R376W, or D1082G) with N-terminal TAP tag fusion were spot tested on rich medium with and without fluconazole at a final concentration of 10 µg/ml along with the strain CBS138 *pdr1*Δ, the wild-type strain CBS138, and the corresponding untagged Pdr1 variants. C. The TAP-tagged strains expressing wild-type Pdr1 or GOF Pdr1 forms were analyzed for Pdr1 and Cdr1 protein levels by western blot analysis using anti-Pdr1 or anti-Cdr1 polyclonal serum.

To co-purify Pdr1-interacting proteins, we carried out standard two-step TAP-purification on native protein extracts from strains expressing TAP-tagged wild-type Pdr1, R376W Pdr1, D1082G Pdr1, or the untagged Pdr1 using IgG- and calmodulin-beads to perform sequential affinity chromatography (Puig *et al*. 2001; Paul *et al*. 2018). The purified proteins were analyzed by multidimensional protein identification technology (MudPIT) (Maccoss *et al*. 2002; Paul *et al*. 2018). Of the several hundred proteins identified in the MudPIT screen, we focused on ones with potential roles in azole resistance, including transcriptional coactivators. Proteins identified in the TAP-tagged Pdr1 samples that were also identified in the untagged wild-type Pdr1 control were considered to have purified independently of Pdr1 and were eliminated from further consideration. (Supplemental Data table 1).

As a first step to examine the relevance of putative Pdr1-interacting factors in azole resistance, we deleted the corresponding open reading frames (ORFs) and screened for changes in azole sensitivity by spot test assay. Our screen included the following genes: *SPO14* (a phospholipase upregulated in azole resistant *C. glabrata* (Vermitsky *et al*. 2006), *CNA1* (the catalytic subunit of calcineurin involved in fluconazole resistance of *C. glabrata* (Chen *et al*. 2012), *SAN1* (a ubiquitin protein ligase upregulated in fluconazole-resistant clinical isolates (Caudle *et al*. 2011), *RIF1* (a telomere-length regulatory protein with roles in subtelomeric silencing (Rosas-Hernandez *et al*. 2008), *FUN30* (a chromatin remodeler of the Snf2 family (Awad *et al*. 2010), *SNF2* (the catalytic subunit of the SWI/SNF chromatin remodeling complex (Laurent *et al*. 1993), *NGG1*/*ADA3* (a subunit of the SAGA coactivator complex (Horiuchi *et al*. 1995);Grant, 1997 #3039}, and *SPT7* (an essential structural subunit of the SAGA coactivator complex (Grant *et al*. 1997; Sterner *et al*. 1999; Wu and Winston 2002). The deletion of *CNA1* had an observable effect on azole sensitivity.

The role of Cna1 in the azole response of *C. glabrata* was characterized by (Vu *et al*. 2023) and will not be discussed here. We observed no significant change in the azole susceptibility of the *spo14Δ*, *san1Δ*, *rif1Δ*, or *fun30Δ* mutants compared to the corresponding wild-type strain (Supplemental Figure 1). Deletion of the SWI/SNF subunit *SNF2* and SAGA complex subunits *SPT7* and *NGG1*/*ADA3* resulted in increased fluconazole sensitivity, indicating a potential function in transcriptional regulation of genes associated with fluconazole resistance (Figure 2). However, the *snf211* and *spt7Δ* mutants exhibited growth and viability defects on rich medium in the absence of azole stress (Figure 2). In order to bypass the generalized deleterious effects of disruption of the SWI/SNF or SAGA coactivator complexes, we chose to examine the effect of acute depletion of coactivator subunits using an auxin-inducible degradation (AID) system (Nishimura *et al*. 2009).

**Figure 2:**
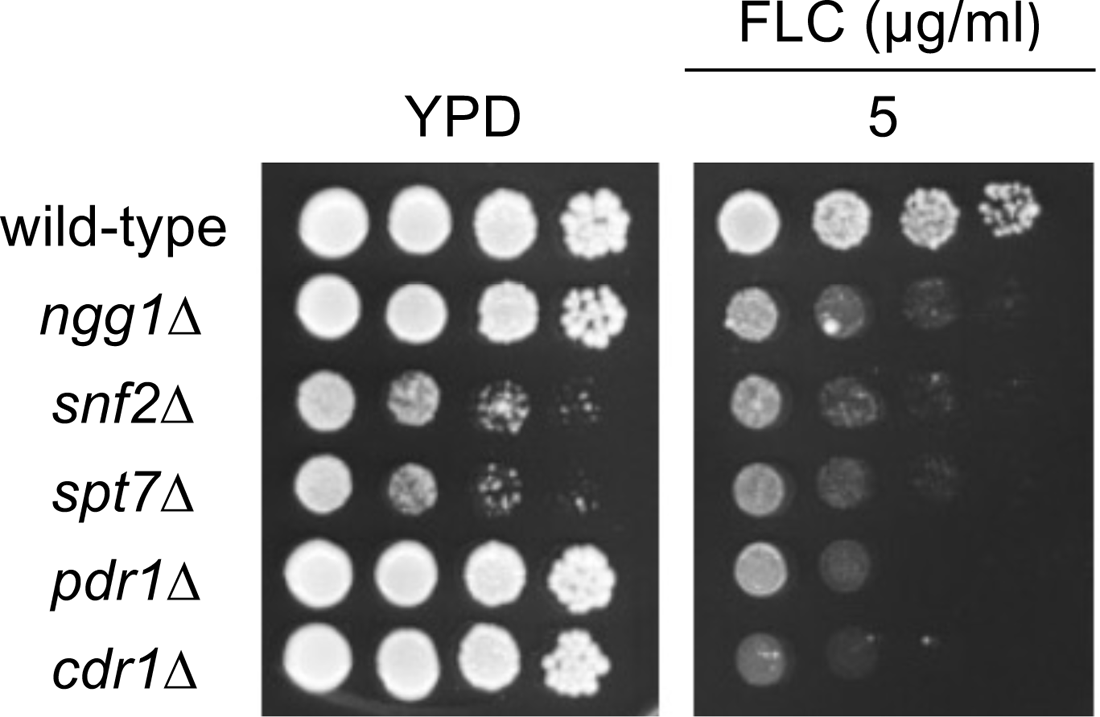
SWI/SNF and SAGA subunit deletions negatively impact azole resistant of *C. glabrata.* The SWI/SNF subunit *SNF2* and the SAGA subunits *SPT7* and *NGG1* were separately deleted from *Candida glabrata* strain CBS138, and the resulting transformants were screened for altered azole sensitivity. Transformants were grown to mid-log phase in rich liquid media (YPD), and serial dilutions of each culture were spotted on YPD agar plates. Where indicated, fluconazole was present at the concentrations shown. Wild-type, *pdr1Δ*, and *cdr1Δ* strains were included in the screen for comparison.

### Conditional depletion of coactivator subunits using an auxin-inducible degradation system

The growth defects of the *snf2*Δ and *spt7*Δ null mutants (Figure 2), prompted us to use an auxin-inducible degradation (AID) strategy (Nishimura *et al*. 2009; Conway and Moye-Rowley 2023; Milholland *et al*. 2023), that allowed for the identification of phenotypes resulting from the acute depletion of individual coactivator subunits. We developed a *C. glabrata* strain modified to constitutively express the *Oryza sativa* auxin receptor F-box protein, OsTir1. Fusions were made at the carboxy-terminus of each coactivator subunit consisting of (i) an auxin-inducible degron (AID) and (ii) a 5X-FLAG motif for monitoring protein levels by western blot. The auxin-inducible degron has a high affinity for the plant hormone auxin and its analogs (Nishimura *et al*. 2009). Upon auxin exposure, the AID domain is triggered to fold and recruit the OsTir1 ligase. As a result of their proximity, the AID-tagged protein is ubiquitinated and subsequently degraded.

To test that each coactivator subunit tolerated the C-terminal AID tag without significantly affecting growth, viability, or azole phenotype, we performed a spot test to compare the AID-tagged coactivator subunit strains to both the wild-type strain as well as the parental strain which constitutively expressed OsTir1 but lacked any AID-tagged proteins. The expression of OsTir1 in the absence of an AID-tagged protein did not affect azole resistance. There was a modest decrease in azole susceptibility in the strain expressing the Snf2-AID but no alteration for any of these AID fusion strains on YPD agar medium (Figure 3A). The AID fusion proteins were well-tolerated and we proceeded with analysis of subunit depletion in the presence of 1-Naphthalene-acetic acid (NAA), a synthetic auxin.

**Figure 3:**
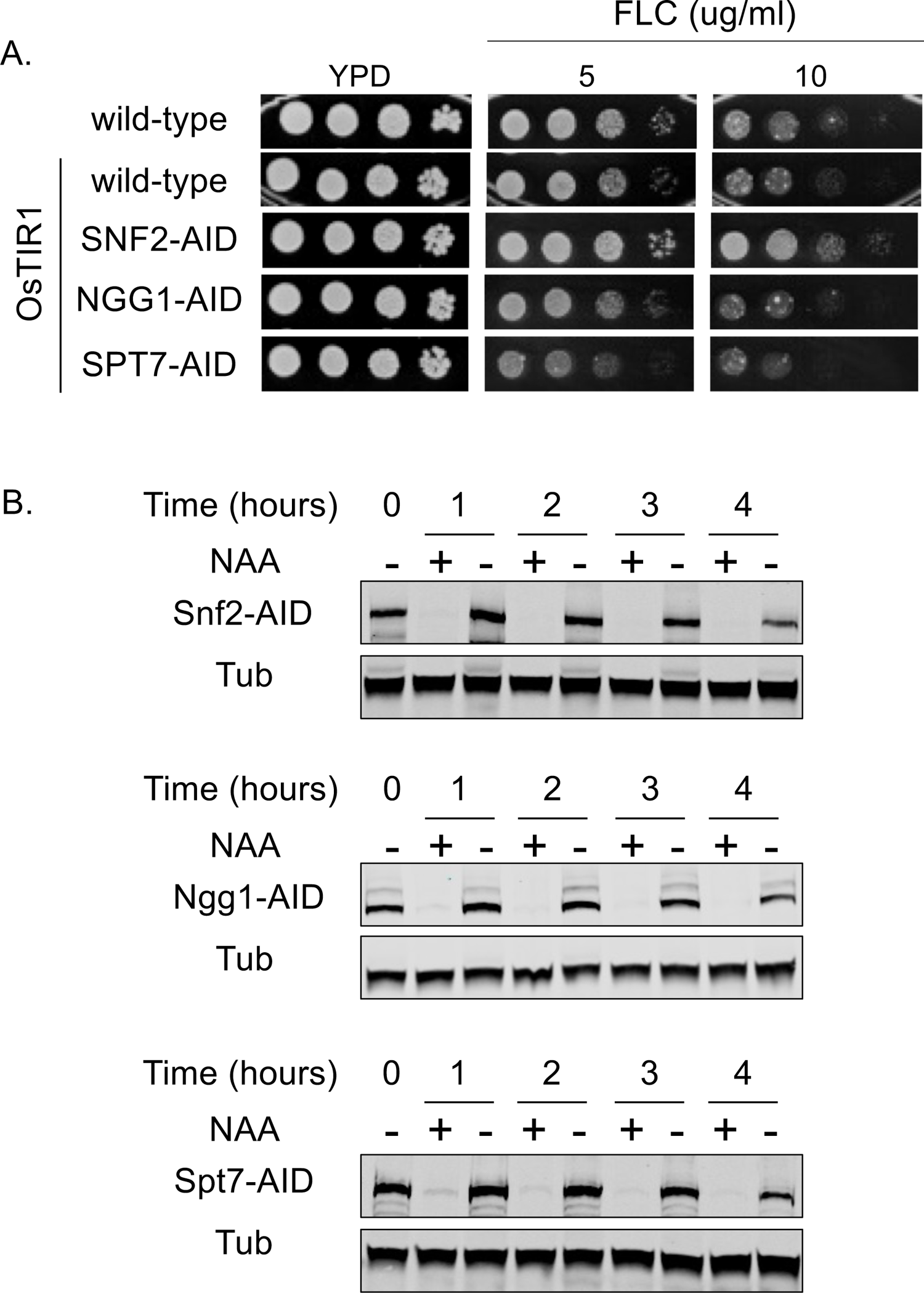
Validation of acute conditional depletion of coactivator subunits using an auxin-inducible degradation strategy. Isogenic *C. glabrata* strains expressing the plant F-box protein, OsTir1, and individual coactivator subunits C-terminally tagged with an auxin-inducible degron (AID) were generated for downstream analysis of the effect of acute coactivator subunit depletion on azole phenotype. A. Spot test assays were performed to ensure that strains expressing OsTir1 and AID-tagged coactivator subunits were tolerated in these strains and did not significantly impact growth or fluconazole (FLC) susceptibility. B. AID-tagged coactivator strains were grown in the presence or absence of 1-Napthaleneacetic acid (NAA, auxin) and western blot was performed to verify protein depletion and validate AID function. AID-tagged strains were grown to mid-log phase in rich medium and split into two cultures, one treated with 0.5 mM NAA (+) and one that remained untreated (-). Samples were taken prior to splitting the cultures and hourly for four hours after addition of NAA. AID-tagged coactivator subunits were detected using an anti-FLAG monoclonal antibody. Tubulin was used as a loading control.

To analyze the timing and efficiency of coactivator depletion in early- and mid-log phase cells cultured in liquid YPD medium, we split early-log cultures of the AID-tagged coactivator strains between two conditions: (i) a control condition consisting only of liquid YPD medium and (ii) a depletion condition of YPD medium containing NAA at a final concentration of 0.5 mM. Samples were removed immediately prior to the splitting of the cultures and subsequently every hour for four hours after exposure to NAA. Using an anti-FLAG monoclonal antibody, the effect of NAA on coactivator subunit degradation was assessed by western blot (Figure 3B). After a one hour incubation with 0.5 mM NAA, all AID-tagged coactivator subunits were undetectable by western blot and remained undetectable during the subsequent hours of NAA exposure, indicating the conditions employed were sufficient for the acute and maintained depletion of the AID-tagged coactivator proteins being examined. Coactivator subunit levels were unaffected when cells were cultured in the absence of NAA.

### Auxin-induced depletion of SWI/SNF or SAGA subunits causes increased azole susceptibility in various Pdr1 backgrounds

To investigate the effect of Snf2, Spt7, or Ngg1 depletion associated with specific Pdr1 variants, we analyzed azole sensitivity of AID-tagged coactivator strains expressing either wild-type-, R376W-, or D1082G-Pdr1 via spot test, in the presence or absence of NAA (Figure 4). For this, we used YPD agar plates containing 1 mM NAA to ensure the maintenance of auxin-induced degradation during the plate incubation time.

**Figure 4:**
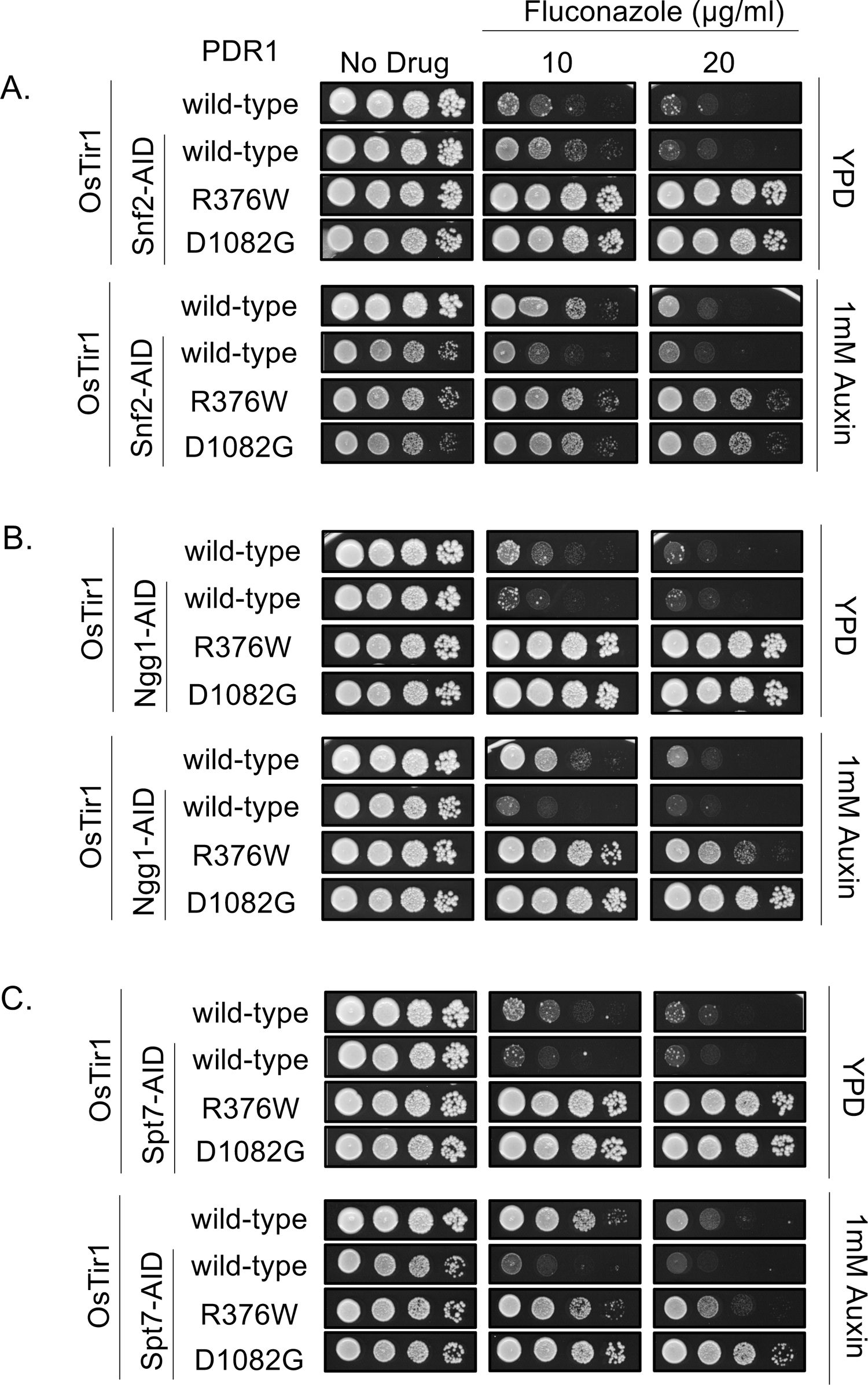
Depletion of SWI/SNF or SAGA subunits increases sensitivity to fluconazole. In a *C. glabrata* strain modified to constitutively express the plant F-box protein OsTir, the SWI/SNF subunit Snf2 (A) or the SAGA subunits Ngg1 (B) or Spt7 (C) were tagged with an auxin-inducible degron (AID) for assessing the effect of conditional coactivator subunit depletion in a wild-type background and two gain-of-function Pdr1 backgrounds, R376W or D1082G. To check whether the AID-tag fusions of coactivator subunits differentially affected fluconazole sensitivity in the three Pdr1 backgrounds, susceptibility to the indicated concentrations of fluconazole (FLC) was assessed on rich medium in the absence of the synthetic plant hormone, 1-Napthaleneacetic acid (NAA, auxin). Degradation phenotypes were assessed on rich medium containing 1mM NAA without FLC and in the presence of FLC at the indicated concentrations.

In the absence of NAA, the presence of the AID-tagged coactivator alleles did not modify the fluconazole susceptibilities of the three *PDR1* alleles; with the strain expressing wild-type Pdr1 being most susceptible to fluconazole and the strains expressing R376W Pdr1 and D1082G Pdr1 being least susceptible (Figure 4A,B,C). The induced degradation of Snf2-AID on YPD agar containing 1 mM NAA resulted in a slight growth defect reminiscent of the *snf2Δ* mutant phenotype (Figure 4A). This was true irrespective of Pdr1 variant.

When strains expressing the Ngg1-AID fusion protein were grown on YPD plates containing 1 mM NAA in the absence of fluconazole, there was no observable growth effect (Figure 4B). Therefore, the acute and maintained depletion of the Ngg1-AID was tolerated, just as the deletion of *NGG1*. When the Ngg1-AID strain expressing wild-type Pdr1 was grown on YPD containing 1mM NAA and varying concentrations of fluconazole, sensitivity to azole increased (Figure 4B). The observed azole phenotype of Ngg1-AID depletion matched that of the *ngg1Δ* strain (Figure 2). Under the conditions tested, Ngg1-AID depletion increased azole sensitivity in cells expressing R376W Pdr1 but had a negligible effect on azole sensitivity in cells expressing D1082G Pdr1 (Figure 4B). In the presence of fluconazole, cells depleted of Spt7-AID exhibited increased azole susceptibility regardless of the Pdr1 form they expressed (Figure 4C). Similar to the effect of Ngg1-AID depletion, Spt7-AID depletion exhibited a greater impact on azole sensitivity in cells expressing R376W Pdr1 than those expressing D1082G Pdr1 (Figure 4C). Together, these data demonstrate a role in azole resistance for the SAGA complex in different Pdr1 backgrounds and indicate that R376W Pdr1 may be more dependent on the presence of a functional SAGA complex than D1082G Pdr1.

### Transcriptional activation by Pdr1 is differentially impacted by depletion of coactivator subunits

To correlate the resistance phenotypes above with gene expression, we carried out mRNA measurements using RT-qPCR analyses on these strains containing auxin-sensitive coactivator fusions. We examined a set of Pdr1 target genes and the *ERG11* gene. *ERG11* encodes the enzymatic target of azole drugs and is transcriptionally induced by fluconazole but is not a target gene of Pdr1 (Whaley *et al*. 2014; Vu and Moye-Rowley 2022). The three Pdr1 target genes we examined were *PDR1* itself, as this gene is positively autoregulated (reviewed in (Moye-Rowley 2019)), along with *CDR1* and *CDR2*, both encoding ABC transporter proteins that have been implicated in fluconazole resistance (Miyazaki *et al*. 1998; Sanglard *et al*. 1999).

For each strain, we removed aliquots prior to exposure to either fluconazole or auxin (pretreatment). These cultures were then divided between two conditions, one containing 0.5 mM NAA and the other lacking NAA. One hour after NAA was introduced to the culture, fluconazole was added to both the NAA-containing culture and the NAA-free culture, to a final concentration of 20 μg/ml. These cultures were incubated for three hours and then levels of *PDR1*, *CDR1*, *CDR2* and *ERG11* mRNA determined using RT-qPCR and appropriate primers.

Using the Snf2-AID strain, auxin-dependent loss of Snf2 led to a reduction in the degree of azole induction of *PDR1* transcription in the presence of wild-type Pdr1 (Figure 5A). Depletion of Snf2 had no effect on the observed level of *PDR1* autoregulation of via either GOF mutant form of Pdr1. *CDR1* transcription was not impacted by Snf2 depletion during this time, irrespective of the Pdr1 derivative present. *CDR2* transcription was elevated in the presence of the D1082G form of Pdr1 prior to FLC treatment. D1082G Pdr1 also drives the highest level of *CDR2* transcription after FLC challenge but this is not impacted by loss of Snf2. Surprisingly, depletion of Snf2 triggered an increase in FLC-induced *CDR2* expression but only in the presence of the wild-type Pdr1. *ERG11* induction by FLC was reduced in Snf2-depleted cells but only modestly.

**Figure 5:**
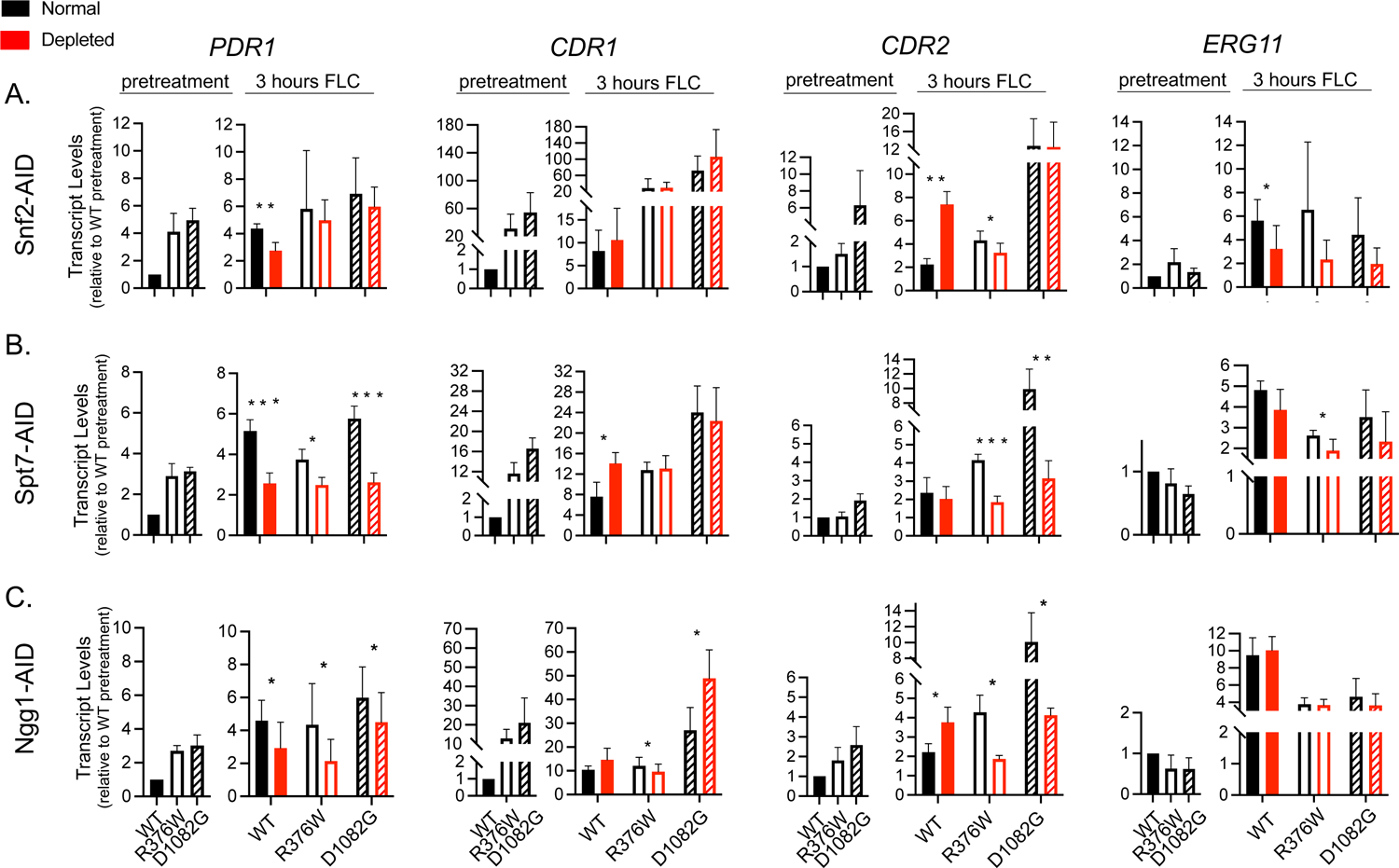
Characterization of SWI/SNF and SAGA subunit contribution to activated transcription of *PDR1, CDR1, CDR2,* and *ERG11.* An auxin-inducible degradation strategy was used to analyze the effects of conditional depletion of the SWI/SNF subunit, Snf2 (A), and the SAGA subunits, Spt7 (B) and Ngg1 (C), on azole-induced transcription of *PDR1, CDR1, CDR2*, and *ERG11* in cells expressing wild-type (solid bars), R376W (empty bars), or D1082G Pdr1 (striped bars). Strains were grown to early-log phase in liquid YPD (pretreatment) and split between two conditions, a normal condition (black) consisting of untreated YPD and in which the subunit of interest was not degraded and a depletion condition (red) consisting of YPD treated with 0.5mM NAA and in which the coactivator subunit of interest was depleted. After the split, cells were cultured to mid-log phase and fluconazole (FLC) was added to both conditions to a final concentration of 20 µg/ml. Samples from both normal and depletion conditions were acquired three hours after the addition of FLC for assessing the effect of depletion on azole-induced transcription. Bar graphs show fold difference in transcript levels of *PDR1, CDR1*, and *ERG11* in all strains relative to the pretreatment levels observed for cells expressing wild-type Pdr1. All measurements represent the averaged result from two independent experiments each with two technical replicates and error bars were calculated as standard deviation.

The two SAGA components will be considered together since their depletion had generally similar effects on expression of Pdr1 target genes. Loss of either Spt7 or Ngg1 led to a reduction in the extent of *PDR1* autoregulation by all forms of Pdr1 although the most significant reduction was seen for wild-type and D1082G forms of Pdr1 upon depletion of Spt7. Ngg1 depletion gave similar effects on *PDR1* transcription but the effect was more modest. FLC-induced *CDR1* transcription was slightly increased in the presence of wild-type Pdr1 while both GOF forms were unaffected upon Spt7 depletion. Depletion of Ngg1 slightly decreased FLC induction of *CDR1* in the presence of the R376W Pdr1 variant while exhibiting an increase in FLC-induced expression when D1082G was present. Loss of either SAGA component reduced FLC induction of *CDR2* expression in the presence of either GOF form of Pdr1 but wild-type Pdr1 drove slightly higher levels of FLC-induced *CDR2* when Ngg1 was depleted. Finally, *ERG11* expression was basically unaffected by depletion of either SAGA component in the presence of FLC and any of the Pdr1 derivatives.

From these data, it appears that *PDR1* transcription is the most sensitive to depletion of any coactivator subunits. We were surprised that the effects on *CDR1* transcription were relatively small, given the clear FLC phenotypes caused by coactivator depletion above (Figure 4). Along with these measurements of the functional consequences caused by coactivator depletion (increased FLC susceptibility and transcriptional alterations), we wanted to assess the potential recruitment of these coactivator proteins to relevant Pdr1 target promoters in the presence and absence of FLC treatment. We used chromatin immunoprecipitation to determine if these coactivator proteins exhibited differential recruitment that was responsive to either changes in Pdr1 variant or the presence of FLC

### Coactivator recruitment to Pdr1 target genes responds to both Pdr1 forms and FLC challenge

We constructed an isogenic series of strains that varied at their *PDR1* locus and contained 3X HA-tagged forms of each of the three coactivator proteins characterized above. These strains contained wild-type, *pdr111*, or the R376W or D1082G GOF forms of Pdr1. Each strain was grown to mid-log phase and then incubated with or without FLC for 1 hour. After this time, chromatin immunoprecipitation (ChIP) was performed and target promoter detected using appropriate PCR primers.

Snf2 was found to be inducibly bound at the *PDR1* promoter (Figure 6A). Both GOF forms of Pdr1 exhibited high, constitutive levels of Snf2 recruitment in the absence of FLC, which was not affected by FLC treatment. We did not assess coactivator binding to the *PDR1* promoter in *pdr1Δ* cells as we were concerned that deletion of the coding sequence of the gene to be assayed for binding might confuse the results of this assay.

**Figure 6:**
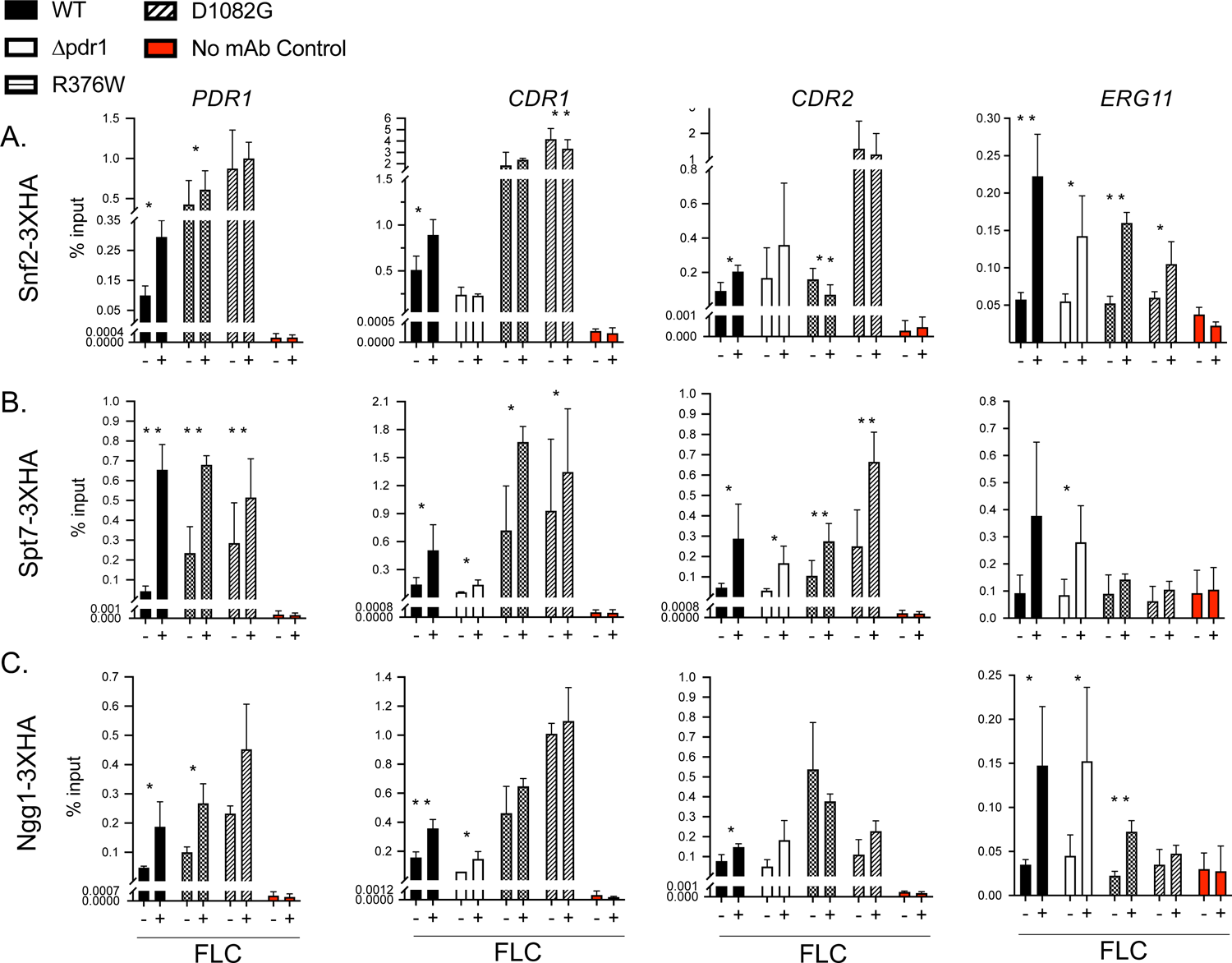
Characterization of the recruitment of SWI/SNF and SAGA subunits to the promoters of *PDR1, CDR1, CDR2,* and *ERG11* in response to azole stress. Chromatin immunoprecipitation (ChIP) of SWI/SNF and SAGA subunits was performed to assay coactivator complex binding at the promoters of the azole-responsive genes, *PDR1, CDR1, CDR2*, and *ERG11*. Strains expressing 3XHA-tagged Snf2 (A, SWI/SNF), Spt7 (B, SAGA), or Ngg1 (C, SAGA) were grown to mid-log phase, treated with fluconazole (FLC) at a final concentration of 20 µg/ml, and grown for one hour in the presence of FLC. Samples were acquired prior to the addition of FLC to the cultures (-) and one hour after the addition of FLC to the cultures (+). Sheared chromatin was prepared from four independent *PDR1* backgrounds for each coactivator subunit analyzed: wild-type *PDR1* (solid bars), *pdr1*Δ (open bars), R376W *PDR1* (checkered bars), and D1082G *PDR1* (striped bars). ChIP reactions were carried out using an anti-HA monoclonal primary antibody (black bars) or in the absence of a primary antibody (red bars) to control for nonspecific immunoprecipitation of genomic DNA. Immunoprecipitated DNA was quantitated by qPCR using primers specific to the *PDR1, CDR1, CDR2*, and *ERG11* promoters, and data are plotted as the percentage of DNA immunoprecipitated relative to total DNA input.

This same pattern of Snf2 binding was seen at the *CDR1* promoter, although more Snf2 was associated with this promoter than *PDR1* (note the different scales of these plots) and there was a clear dependence on the presence of Pdr1 to recruit Snf2 to this promoter, especially in the case of the GOF forms of Pdr1. *CDR2* showed no significant binding of Snf2 except in the presence of the D1082G GOF variant of Pdr1. Finally, Snf2 recruitment to *ERG11* was induced in the presence of FLC and not significantly responsive to the *PDR1* variant present, findings consistent with the fact that Pdr1 does not control *ERG11* transcription.

The recruitment of the SAGA components was similar between Spt7 (Figure 6B) and Ngg1 (Figure 6C) but showed some interesting differences. Spt7 binding to the *PDR1* promoter was highly responsive to FLC challenge but also to the presence of Pdr1. Both GOF forms of Pdr1 exhibited Spt7 recruitment in the absence of FLC treatment that could be further induced by treatment with this antifungal drug. The association of Ngg1 to the *PDR1* promoter was similar but of a reduced magnitude.

The degree of association of Spt7 with the *CDR1* promoter showed a reduced response to FLC treatment compared to *PDR1* (∼2-fold compared to >10-fold) but was highly dependent on the presence of Pdr1 and strongly responsive to the presence of both GOF forms of Pdr1. Ngg1 showed this same behavior.

Spt7 engagement with the *CDR2* promoter was induced by FLC exposure but this appeared primarily Pdr1 independent. The exception to this behavior was seen in the presence of the D1082G Pdr1 variant that showed the highest level of Spt7 recruitment in the presence of FLC. Strikingly, Ngg1 recruitment to *CDR2* was quite modest, with the exception of the R376W Pdr1 protein that showed the highest level of Ngg1 association with *CDR2,* irrespective of the presence of FLC. This non-identical SAGA complex binding to the *CDR2* promoter illustrated the difference in behaviors between the D1082G and R376W forms of Pdr1 with this differential recruitment of Spt7 and Ngg1.

Finally, *ERG11* promoter association levels of both Spt7 and Ngg1 were increased by FLC treatment. Recruitment of these coactivators was decreased by the presence of the GOF forms of Pdr1 although a degree of FLC induction was maintained for Ngg1 recruitment in the presence of the R376W *PDR1* allele. This reduced *ERG11* association is likely due to the well-described reduction in FLC susceptibility caused by the presence of these GOF Pdr1 mutants (reviewed in (Whaley and Rogers 2016)).

### Overlapping roles of Snf2 and Spt7 during FLC-induced gene induction

The data above provide strong evidence that both Snf2 and Spt7/Ngg1 can be found at Pdr1-responsive promoters and contribute to the regulation of gene expression there. The presence of both these factors led us to investigate if loss of one transcriptional coactivator could be suppressed by the action of the other. This functional redundancy has been seen for Snf2 and the histone acetyltransferase Gcn5 in *S. cerevisiae* in earlier work (Sudarsanam *et al*. 1999) as well as for Snf2 and the histone chaperone Asf1 (Gkikopoulos *et al*. 2009).

To determine if Snf2 and Spt7 might be acting in a redundant manner at Pdr1-regulated genes, we constructed a strain containing auxin-depletable forms of both of these coactivator proteins. This strain was grown to early log phase, split into two aliquots in the presence or absence of auxin and then challenged with FLC for 3 hrs. We then measured RNA levels for *PDR1, CDR1, CDR2* and *ERG11* using RT-qPCR as before (Figure 7A).

**Figure 7:**
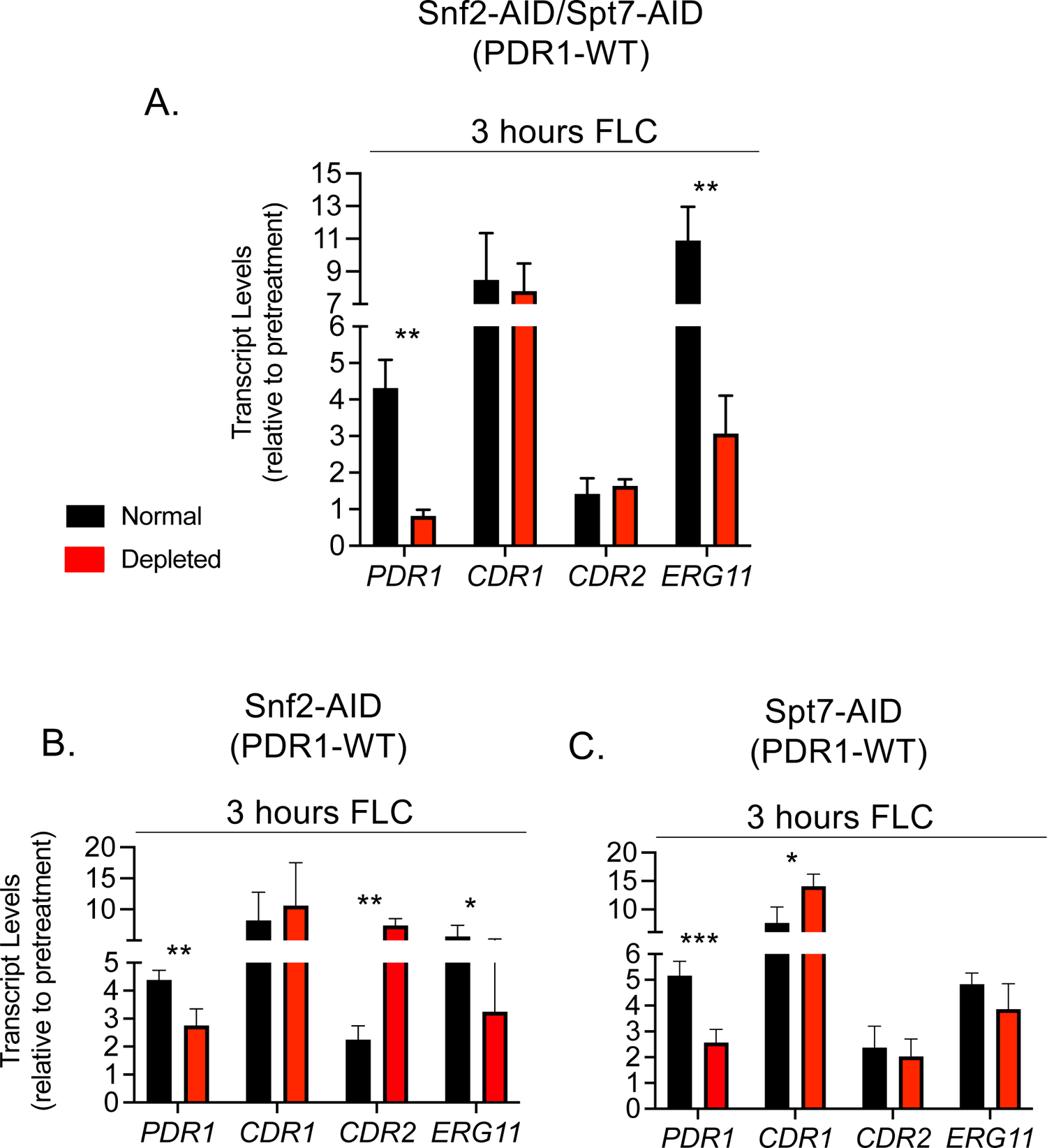
Fluconazole-induced gene expression after simultaneous disruption of both SWI/SNF and SAGA. Auxin-induced degradation was used to simultaneously disrupt the SWI/SNF and SAGA coactivator complexes to determine their cumulative contribution to activated transcription of *PDR1, CDR1, CDR2*, and *ERG11*. The experiment was performed as previously described in order to compare untreated (normal) and auxin-treated (depleted) cultures. Samples were acquired for qPCR analysis prior to the splitting of the early-log phase culture (pretreatment) and three hours after the addition of FLC (i.e. four hours after auxin treatment). Data is presented as the fold difference between pretreatment transcript levels and FLC-induced transcript levels for both the normal (black) and depleted (red) conditions. Results for individual depletion of Snf2 (B) or Spt7 (C) are also shown for comparison.

Auxin depletion of both Snf2 and Spt7 blocked FLC induction of *PDR1* and lowered FLC induction of *ERG11* from 11-fold to roughly 3-fold. *CDR1* and *CDR2* were not affected by this double depletion of both Snf2 and Spt7. Loss of either coactivator alone (shown in Figure 7B and 7C) still retained more than 2-fold induction by FLC for *PDR1*. These data argue that Snf2 and Spt7 are both critical for the FLC induction of *PDR1* as their simultaneous loss led to elimination of any detectable FLC induction of this gene.

## Discussion

The *PDR1* gene has been extensively documented to be the main site of mutations leading to reduced FLC susceptibility in *C. glabrata*. Many groups have described substitution mutations that cause Pdr1 to exhibit elevated, constitutive activation of downstream target genes, like the ABC transporter-encoding locus *CDR1* (Vermitsky and Edlind 2004; Tsai *et al*. 2006; Vermitsky *et al*. 2006; Ferrari *et al*. 2009; KhakhinA *et al*. 2018; Salazar *et al*. 2022). However, there is little known about the molecular basis underpinning the altered activity states of Pdr1 or how it leads to enhanced downstream gene expression. We performed a copurification screen which indicated Snf2, the catalytic subunit of the SWI/SNF chromatin remodeling complex, as well as Ngg1/Ada3 and Spt7, two subunits from the SAGA coactivator complex, interact with Pdr1 variants. The SWI/SNF and SAGA coactivator complexes are highly conserved among eukaryotes and have been demonstrated to function in gene activation via chromatin remodeling and the modification of histones, respectively (Rando and Winston 2012).

Deletion of the *SNF2, SPT7*, or *NGG1* genes increased the azole sensitivity of *C. glabrata*, findings consistent with those of others demonstrating increased azole susceptibility in *C. glabrata snf2*Δ and *ngg1*Δ strains (Nikolov *et al*. 2022; Lin *et al*. 2023). We observed that permanent disruption of SWI/SNF or SAGA function via deletion of *SNF2* or *SPT7*, respectively, resulted in systemic consequences for *C. glabrata* that manifested as slowed growth and decreased viability. To limit defects associated with permanent loss of essential coactivator subunits, we chose an auxin-inducible protein depletion strategy to examine the effect of coactivator subunit loss.

The conditional nature of the depletion strategy allows cells to maintain functional coactivator complexes until acute depletion of the subunit(s) of interest, thereby reducing the more general consequences triggered by chronic elimination of complex function in deletion mutants.

Another feature of the rapid auxin depletion strategy is worthy of consideration. The depletion of these chromatin-modifying proteins, rather than their chronic absence, is likely to reveal more direct functions of these factors in the transcriptional response to azole drugs. For example, experiments with auxin degradation of SAGA core module subunits (i.e. Spt3 or Spt7) in *S. cerevisiae* showed that while these factors were rapidly eliminated from the cell, modifications that depend on these proteins were slower to be lost (Donczew *et al*. 2020). Thus, defects that present rapidly after loss of a coactivator protein are expected to pinpoint direct effects such as SAGA-dependent TATA-binding protein loading as has been described (reviewed in (Timmers 2021)). Thus, the effects of acute coactivator subunit depletion, as reported here, reflect functionality of the coactivator at the onset of azole exposure.

The SWI/SNF chromatin remodeling complex has been shown to function in virulence and the transcriptional response to environmental stress in fungal pathogens (reviewed in (Balachandra and Ghosh 2022)). We observed that the acute disruption of SWI/SNF function via conditional depletion of Snf2 had an inhibitory effect on FLC-activated transcription of *PDR1* in cells expressing wild-type Pdr1 but not in cells expressing either R376W or D1082G Pdr1 (Figure 5A). Coupled with our observation that GOF Pdr1 variants have already recruited Snf2 to the *PDR1* promoter while the wild-type protein must be activated by FLC for this recruitment to occur (Figure 7A), we suggest that an early step in Pdr1 autoregulation is likely to be this accumulation of SWI/SNF at the *PDR1* promoter.

Evidence for promoter-specific effects of Pdr1 comes from the comparison of *PDR1* autoregulation (described above) and FLC-activated transcription of the Pdr1-responsive genes *CDR1* and *CDR2*. Both of these ABC transporter-encoding loci failed to exhibit a significant dependence on SWI/SNF function during FLC induction of transcription by wild-type Pdr1. While our findings broadly agree with those reported earlier for SWI/SNF playing a role in Pdr1-driven gene activation (Nikolov *et al*. 2022), the conditional depletion strategy employed here sheds light on promoter-specific roles for SWI/SNF at Pdr1-regulated loci and how they differ. The depletion of Snf2 caused a reduction in the expression of *ERG11*, irrespective of the *PDR1* allele tested, which illustrates the global nature of Snf2 involvement in promoter activation.

Previous studies of the SAGA coactivator complex have provided evidence for its multiple functions in transcriptional activation, including histone acetylation and deubiquitylation and its direct interactions with transcription factors (e.g. TATA-binding protein) (reviewed in (Baker and Grant 2007)). Here, we have shown that disruption of SAGA complex function via deletion or conditional depletion of the SAGA subunits Spt7 or Ngg1 increased azole sensitivity of the corresponding cells. This finding is in agreement with previous observations that deletion of SAGA histone acetyltransferase module subunits (i.e. *GCN5, ADA2, NGG1*) negatively impacts azole resistance in *C. glabrata* (Lin *et al*. 2023). Using the auxin-inducible protein depletion strategy to examine the effect of acute loss of either Spt7 or Ngg1 on Pdr1-dependent gene activation, we demonstrated that disruption of SAGA diminished FLC-activated *PDR1* transcription at the onset of azole challenge. Unlike the observed Pdr1 variant-dependent effect of SWI/SNF, acute depletion of Spt7 or Ngg1 inhibited azole-induced *PDR1* transcription in cells expressing either wild-type Pdr1 or GOF Pdr1 forms.

Furthermore, transcription of *ERG11* was relatively unaffected by acute depletion of either Spt7 or Ngg1 in all strain backgrounds tested, illustrating the greater reliance on SAGA function at Pdr1-dependent promoters. Consistent with a key role in Pdr1-dependent gene regulation, SAGA complex subunits exhibited azole-induced recruitment to the *PDR1* and *CDR1* promoters independent of the Pdr1 form expressed. We also detected a Pdr1 GOF variant-specific effect caused by depletion of Ngg1 (Figure 5C). While depletion of Ngg1 caused a reduction in R376W Pdr1-driven *CDR1* expression, *CDR1* expression in the presence of the D1082G form of Pdr1 was not affected. This observation fits well with our previous genetic data that the mechanisms used by different GOF forms of Pdr1 are not identical and may exhibit different coactivator dependences (Simonicova and Moye-Rowley 2020). Taken together, these data suggest a requirement for SAGA function in both the transition from basal transcription of *PDR1* to fluconazole-induced transcription of *PDR1*, as well as the maintenance of Pdr1 GOF-driven transcription of *PDR1*.

In these analyses of the effects of acute disruption of SWI/SNF or SAGA, we observed that rapid coactivator subunit depletion caused a typically 2-fold or less reduction of *PDR1* and *CDR1* transcription. We wondered if this relatively modest reduction might be due to a functional suppression by the remaining coactivator complex. Cooperative action between SAGA and SWI/SNF has been described in the transcriptional activation of cell wall integrity genes as well as genes required for non-fermentative growth in *S. cerevisiae* (Biddick *et al*. 2008; Sanz *et al*. 2016; Menezes *et al*. 2017). Therefore, we considered that the effect of disrupting one coactivator complex may be buffered by increased or overlapping functions of another complex. To determine whether SWI/SNF and SAGA function cooperatively in FLC-induced gene expression, we analyzed the effect of simultaneously disrupting the complexes by depleting cells of both Snf2 and Spt7. The conditional depletion of both Snf2 and Spt7 resulted in a complete inhibition of FLC-activated *PDR1* and *ERG11* transcription (Figure 7A), indicating that SWI/SNF and SAGA are both required for activated transcription of *PDR1* and *ERG11* and suggesting a likely overlapping nature to their function in azole-induced gene expression. Under the conditions tested the simultaneous disruption of SWI/SNF and SAGA function did not affect FLC-activated transcription of *CDR1* or *CDR2*, further illustrating promoter-specific function of general coactivators and the complex regulatory mechanisms evolved for genes essential to survival under azole stress. Identification of general coactivators required for Pdr1-driven gene activation advances the understanding of mechanisms underlying hyper-resistant phenotypes observed among clinical isolates and contributes to the advancement of therapeutic strategies.

## Data availability statement

Strains and plasmids are available upon request. The authors affirm that all data necessary for confirming the conclusions of the article are present within the article, figures, and tables.

## Acknowledgements

We thank Dr. Damian Krysan for helpful comments on this work. This work was supported by NIH AI152494.

---

**Supplementary Table 1:** A list of co-purified proteins identified by MudPIT analysis. The purified TAP-Pdr1 protein versions (wild type Pdr1, R376W and D1082G) were analyzed by MudPIT technology to identify Pdr1-interacting partners. MS/MS spectral data of each peptide were searched against a *C. glabrata* protein database using Sequest (Yates *et al*. 1995)) and the resulting identifications were collated and filtered using Scaffold (https://www.proteomesoftware.com). The abundance of a peptide is measured as the number of peptide-to-spectrum matching (PSM) events.

**Supplemental Figure 1:**
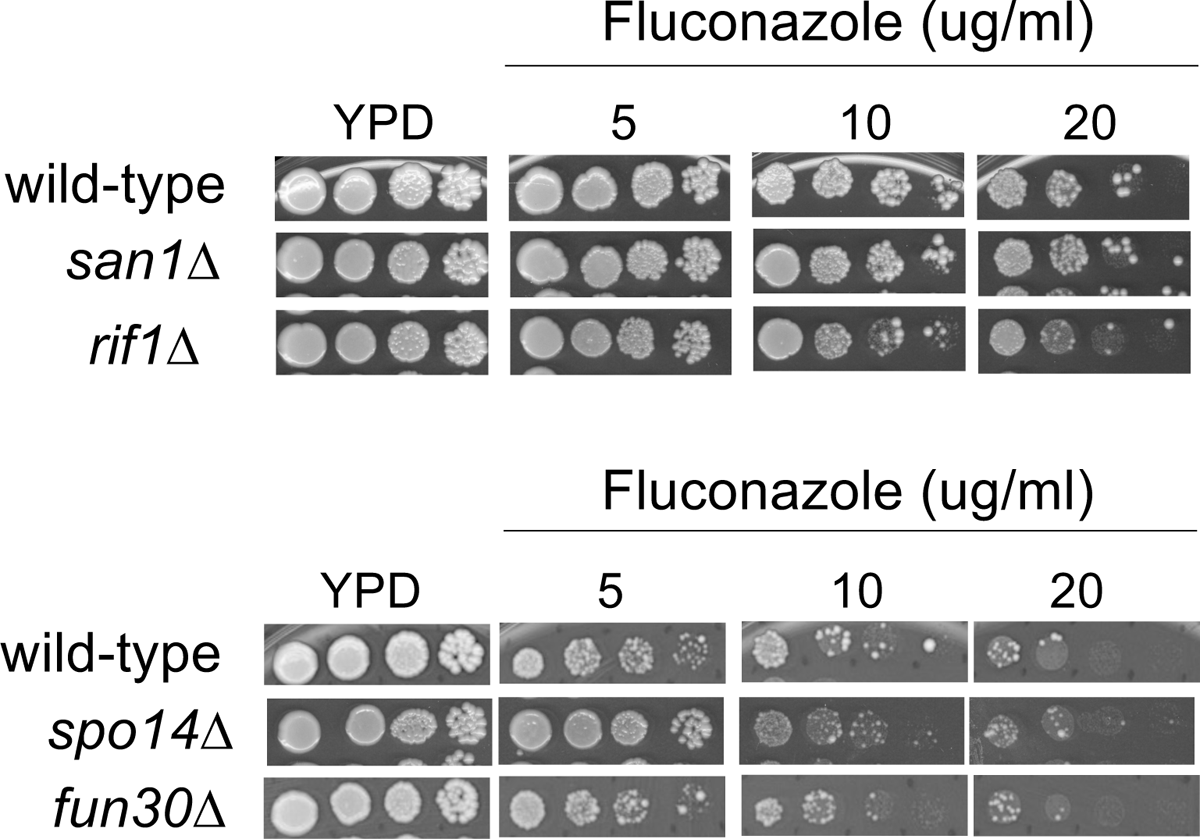
Azole susceptibility of *C. glabrata* mutants from MudPIT screen. Genes identified in the screen for proteins that interact with Pdr1 were deleted from *Candida glabrata* strain CBS138, and the resulting transformants were screened for altered azole sensitivity by spot test. The *san1*Δ, *rif1*Δ, *spo14*Δ, and *fun30*Δ mutants were compared to the wild-type parental strain as a control.

## References

Awad, S., D. Ryan, P. Prochasson, T. Owen-Hughes and A. H. Hassan, 2010 The Snf2 homolog Fun30 acts as a homodimeric ATP-dependent chromatin-remodeling enzyme. J Biol Chem 285: 9477–9484.

Baker, S. P., and P. A. Grant, 2007 The SAGA continues: expanding the cellular role of a transcriptional co-activator complex. Oncogene 26: 5329–5340.

Balachandra, V. K., and S. K. Ghosh, 2022 Emerging roles of SWI/SNF remodelers in fungal pathogens. Curr Genet 68: 195–206.

Biddick, R. K., G. L. Law, K. K. Chin and E. T. Young, 2008 The transcriptional coactivators SAGA, SWI/SNF, and mediator make distinct contributions to activation of glucose-repressed genes. J Biol Chem 283: 33101–33109.

Casamassimi, A., and C. Napoli, 2007 Mediator complexes and eukaryotic transcription regulation: an overview. Biochemie 89: 1439–1446.

Caudle, K. E., K. S. Barker, N. P. Wiederhold, L. Xu, R. Homayouni et al., 2011 Genomewide expression profile analysis of the Candida glabrata Pdr1 regulon. Eukaryot Cell 10: 373–383.

Chen, Y. L., J. H. Konieczka, D. J. Springer, S. E. Bowen, J. Zhang et al., 2012 Convergent Evolution of Calcineurin Pathway Roles in Thermotolerance and Virulence in Candida glabrata. G3 (Bethesda) 2: 675–691.

Conway, T. P., and W. S. Moye-Rowley, 2023 Conditional Protein Depletion in the Analysis of Antifungal Drug Resistance in Candida glabrata. Methods Mol Biol 2658: 191–200.

Donczew, R., L. Warfield, D. Pacheco, A. Erijman and S. Hahn, 2020 Two roles for the yeast transcription coactivator SAGA and a set of genes redundantly regulated by TFIID and SAGA. Elife 9.

Ferrari, S., F. Ischer, D. Calabrese, B. Posteraro, M. Sanguinetti et al., 2009 Gain of function mutations in CgPDR1 of Candida glabrata not only mediate antifungal resistance but also enhance virulence. PLoS Pathog 5: e1000268.

Fisher, M. C., and D. W. Denning, 2023 The WHO fungal priority pathogens list as a game-changer. Nat Rev Microbiol 21: 211–212.

Gkikopoulos, T., K. M. Havas, H. Dewar and T. Owen-Hughes, 2009 SWI/SNF and Asf1p cooperate to displace histones during induction of the saccharomyces cerevisiae HO promoter. Mol Cell Biol 29: 4057–4066.

Grahl, N., E. G. Demers, A. W. Crocker and D. A. Hogan, 2017 Use of RNA-Protein Complexes for Genome Editing in Non-albicans Candida Species. mSphere 2.

Grant, P. A., L. Duggan, J. Cote, S. M. Roberts, J. E. Brownell et al., 1997 Yeast Gcn5 functions in two multisubunit complexes to acetylate nucleosomal histones: characterization of an Ada complex and the SAGA (Spt/Ada) complex. Genes Dev 11: 1640–1650.

Healey, K. R., and D. S. Perlin, 2018 Fungal Resistance to Echinocandins and the MDR Phenomenon in Candida glabrata. J Fungi (Basel) 4.

Horiuchi, J., N. Silverman, G. A. Marcus and L. Guarente, 1995 ADA3, a putative transcriptional adaptor, consists of two separable domains and interacts with ADA2 and GCN5 in a trimeric complex. Mol Cell Biol 15: 1203–1209.

Khakhina, S., L. Simonicova and W. S. Moye-Rowley, 2018 Positive autoregulation and repression of transactivation are key regulatory features of the Candida glabrata Pdr1 transcription factor. Mol Microbiol 107: 747–764.

Kornberg, R. D., 2005 Mediator and the mechanism of transcriptional activation. Trends Biochem Sci 30: 235–239.

Krysan, D. J., 2017 The unmet clinical need of novel antifungal drugs. Virulence 8: 135–137.

Laurent, B. C., I. Treich and M. Carlson, 1993 The yeast SNF2/SWI2 protein has DNA-stimulated ATPase activity required for transcriptional activation. Genes Dev 7: 583–591.

Lin, C. J., S. Y. Yang, L. H. Hsu, S. J. Yu and Y. L. Chen, 2023 The Gcn5-Ada2-Ada3 histone acetyltransferase module has divergent roles in pathogenesis of Candida glabrata. Med Mycol 61.

Livak, K. J., and T. D. Schmittgen, 2001 Analysis of relative gene expression data using real-time quantitative PCR and the 2(-Delta Delta C(T)) Method. Methods 25: 402–408.

MacCoss, M. J., W. H. McDonald, A. Saraf, R. Sadygov, J. M. Clark et al., 2002 Shotgun identification of protein modifications from protein complexes and lens tissue. Proc Natl Acad Sci U S A 99: 7900–7905.

Malik, S., and R. G. Roeder, 2023 Regulation of the RNA polymerase II pre-initiation complex by its associated coactivators. Nat Rev Genet 24: 767–782.

Menezes, R. A., C. Pimentel, A. R. Silva, C. Amaral, J. Merhej et al., 2017 Mediator, SWI/SNF and SAGA complexes regulate Yap8-dependent transcriptional activation of ACR2 in response to arsenate. Biochim Biophys Acta Gene Regul Mech 1860: 472–481.

Milholland, K. L., J. B. Gregor, S. Hoda, S. Piriz-Antunez, E. Duenas-Santero et al., 2023 Rapid, efficient auxin-inducible protein degradation in Candida pathogens. mSphere 8: e0028323.

Miyazaki, H., Y. Miyazaki, A. Geber, T. Parkinson, C. Hitchcock et al., 1998 Fluconazole resistance associated with drug efflux and increased transcription of a drug transporter gene, PDH1, in Candida glabrata. Antimicrob. Agents Chemother. 42: 1695–1701.

Moye-Rowley, W. S., 2019 Multiple interfaces control activity of the Candida glabrata Pdr1 transcription factor mediating azole drug resistance. Curr Genet 65: 103–108.

Nett, J. E., and D. R. Andes, 2016 Antifungal Agents: Spectrum of Activity, Pharmacology, and Clinical Indications. Infect Dis Clin North Am 30: 51–83.

Nikolov, V. N., D. Malavia and T. Kubota, 2022 SWI/SNF and the histone chaperone Rtt106 drive expression of the Pleiotropic Drug Resistance network genes. Nat Commun 13: 1968.

Nishimura, K., T. Fukagawa, H. Takisawa, T. Kakimoto and M. Kanemaki, 2009 An auxin-based degron system for the rapid depletion of proteins in nonplant cells. Nat Methods 6: 917–922.

Paul, S., T. B. Bair and W. S. Moye-Rowley, 2014 Identification of Genomic Binding Sites for Candida glabrata Pdr1 Transcription Factor in Wild-Type and rho0 Cells. Antimicrob Agents Chemother 58: 6904–6912.

Paul, S., W. H. McDonald and W. S. Moye-Rowley, 2018 Negative regulation of Candida glabrata Pdr1 by the deubiquitinase subunit Bre5 occurs in a ubiquitin independent manner. Mol Microbiol 110: 309–323.

Paul, S., and W. S. Moye-Rowley, 2014 Multidrug resistance in fungi: regulation of transporter-encoding gene expression. Front Physiol 5: 143.

Paul, S., J. A. Schmidt and W. S. Moye-Rowley, 2011 Regulation of the CgPdr1 transcription factor from the pathogen Candida glabrata. Eukaryot Cell 10: 187–197.

Puig, O., F. Caspary, G. Rigaut, B. Rutz, E. Bouveret et al., 2001 The tandem affinity purification (TAP) method: a general procedure of protein complex purification. Methods 24: 218–229.

Rando, O. J., and F. Winston, 2012 Chromatin and transcription in yeast. Genetics 190: 351–387.

Rigaut, G., A. Shevchenko, B. Rutz, M. Wilm, M. Mann et al., 1999 A generic protein purification method for protein complex characterization and proteome exploration. Nat Biotechnol 17: 1030–1032.

Rosas-Hernandez, L. L., A. Juarez-Reyes, O. E. Arroyo-Helguera, A. De Las Penas, S. J. Pan et al., 2008 yKu70/yKu80 and Rif1 regulate silencing differentially at telomeres in Candida glabrata. Eukaryot Cell 7: 2168–2178.

Sagatova, A. A., M. V. Keniya, R. K. Wilson, B. C. Monk and J. D. Tyndall, 2015 Structural Insights into Binding of the Antifungal Drug Fluconazole to Saccharomyces cerevisiae Lanosterol 14alpha-Demethylase. Antimicrob Agents Chemother 59: 4982–4989.

Salazar, S. B., M. J. F. Pinheiro, D. Sotti-Novais, A. R. Soares, M. M. Lopes et al., 2022 Disclosing azole resistance mechanisms in resistant Candida glabrata strains encoding wild-type or gain-of-function CgPDR1 alleles through comparative genomics and transcriptomics. G3 (Bethesda) 12.

Sanglard, D., F. Ischer and J. Bille, 2001 Role of ATP-binding cassette transporter gene in high-frequency acquisition of resistance to azole antifungals in Candida glabrata. Antimicrob. Agents Chemother. 45: 1174–1183.

Sanglard, D., F. Ischer, D. Calabrese, P. A. Majcherczyk and J. Bille, 1999 The ATP binding cassette transporter gene CgCDR1 from Candida glabrata is involved in the resistance of clinical isolates to azole antifungal agents. Antimicrob Agents Chemother 43: 2753–2765.

Sanz, A. B., R. Garcia, J. M. Rodriguez-Pena, C. Nombela and J. Arroyo, 2016 Cooperation between SAGA and SWI/SNF complexes is required for efficient transcriptional responses regulated by the yeast MAPK Slt2. Nucleic Acids Res 44: 7159–7172.

Simonicova, L., and W. S. Moye-Rowley, 2020 Functional information from clinically-derived drug resistant forms of the Candida glabrata Pdr1 transcription factor. PLoS Genet 16: e1009005.

Sterner, D. E., P. A. Grant, S. M. Roberts, L. J. Duggan, R. Belotserkovskaya et al., 1999 Functional organization of the yeast SAGA complex: distinct components involved in structural integrity, nucleosome acetylation, and TATA-binding protein interaction. Mol Cell Biol 19: 86–98.

Sudarsanam, P., Y. Cao, L. Wu, B. C. Laurent and F. Winston, 1999 The nucleosome remodeling complex, Snf/Swi, is required for the maintenance of transcription in vivo and is partially redundant with the histone acetyltransferase, Gcn5. EMBO J 18: 3101–3106.

Thakur, J. K., H. Arthanari, F. Yang, S.-J. Pan, X. Fan et al., 2008 A nuclear receptor-like pathway regulating multidrug resistance in fungi. Nature 452: 604–609.

Timmers, H. T. M., 2021 SAGA and TFIID: Friends of TBP drifting apart. Biochim Biophys Acta Gene Regul Mech 1864: 194604.

Tsai, H. F., A. A. Krol, K. E. Sarti and J. E. Bennett, 2006 Candida glabrata PDR1, a transcriptional regulator of a pleiotropic drug resistance network, mediates azole resistance in clinical isolates and petite mutants. Antimicrob Agents Chemother 50: 1384–1392.

Tsai, H. F., L. R. Sammons, X. Zhang, S. D. Suffis, Q. Su et al., 2010 Microarray and molecular analyses of the azole resistance mechanism in Candida glabrata oropharyngeal isolates. Antimicrob Agents Chemother 54: 3308–3317.

Vermitsky, J. P., K. D. Earhart, W. L. Smith, R. Homayouni, T. D. Edlind et al., 2006 Pdr1 regulates multidrug resistance in Candida glabrata: gene disruption and genome-wide expression studies. Mol Microbiol 61: 704–722.

Vermitsky, J. P., and T. D. Edlind, 2004 Azole resistance in Candida glabrata: coordinate upregulation of multidrug transporters and evidence for a Pdr1-like transcription factor. Antimicrob Agents Chemother 48: 3773–3781.

Vu, B. G., and W. S. Moye-Rowley, 2022 Azole-Resistant Alleles of ERG11 in Candida glabrata Trigger Activation of the Pdr1 and Upc2A Transcription Factors. Antimicrob Agents Chemother 66: e0209821.

Vu, B. G., L. Simonicova and W. S. Moye-Rowley, 2023 Calcineurin is required for Candida glabrata Pdr1 transcriptional activation. mBio 14: e0241623.

Vu, B. G., M. A. Stamnes, Y. Li, P. D. Rogers and W. S. Moye-Rowley, 2021 The Candida glabrata Upc2A transcription factor is a global regulator of antifungal drug resistance pathways. PLoS Genet 17: e1009582.

Vu, B. G., G. H. Thomas and W. S. Moye-Rowley, 2019 Evidence that Ergosterol Biosynthesis Modulates Activity of the Pdr1 Transcription Factor in Candida glabrata. MBio 10.

Whaley, S. G., K. E. Caudle, J. P. Vermitsky, S. G. Chadwick, G. Toner et al., 2014 UPC2A is required for high-level azole antifungal resistance in Candida glabrata. Antimicrob Agents Chemother 58: 4543–4554.

Whaley, S. G., and P. D. Rogers, 2016 Azole Resistance in Candida glabrata. Curr Infect Dis Rep 18: 41.

Wu, P. Y., and F. Winston, 2002 Analysis of Spt7 function in the Saccharomyces cerevisiae SAGA coactivator complex. Mol Cell Biol 22: 5367–5379.

Yates, J. R, 3rd., J. K. Eng, A. L. McCormack and D. Schieltz, 1995 Method to correlate tandem mass spectra of modified peptides to amino acid sequences in the protein database. Anal Chem 67: 1426–1436.

